# *In Silico* Analysis Reveals a Shared Immune Signature in *CASP8*-Mutated Carcinomas with Varying Correlations to Prognosis

**DOI:** 10.1101/305755

**Authors:** Yashoda Ghanekar, Subhashini Sadasivam

**Affiliations:** DeepSeeq Bioinformatics, Bangalore 560 064, INDIA; Institute for Stem Cell Biology and Regenerative Medicine (inStem), National Centre for Biological Sciences, UAS-GKVK campus, Yelahanka, Bangalore 560 065, INDIA

**Keywords:** *CASP8*, HNSC, UCEC, Inflammation, Necroptosis, HNSC-TCGA, Neutrophils, IL33

## Abstract

**Background:** Sequencing studies across multiple cancers continue to reveal the spectrum of mutations and genes involved in the pathobiology of these cancers. Exome sequencing of oral cancers, a subset of Head and Neck Squamous cell Carcinomas (HNSCs) common among tobacco-chewing populations, revealed that ~34% of the affected patients harbor mutations in the *CASP8* gene. Uterine Corpus Endometrial Carcinoma (UCEC) is another cancer type where about 10% cases harbor *CASP8* mutations. Caspase-8, the protease encoded by *CASP8* gene, plays a dual role in programmed cell death, which in turn has an important role in tumor cell death and drug resistance. CASP8 is a protease required for the extrinsic pathway of apoptosis and is also a negative regulator of necroptosis. Using bioinformatics approaches to mine data in The Cancer Genome Atlas, we compared the molecular features and survival of these carcinomas with and without *CASP8* mutations.

**Results:** Our *in silico* analyses showed that HNSCs with *CASP8* mutations displayed a prominent signature of genes involved in immune response and inflammation, and were rich in immune cell infiltrates. However, in contrast to Human Papilloma Virus-positive HNSCs, a subtype that exhibits high immune cell infiltration and better overall survival, HNSC patients with mutant-*CASP8* tumors did not display any survival advantage. A similar bioinformatic analyses in UCECs revealed that while UCECs with *CASP8* mutations also displayed an immune signature, they had better overall survival, in contrast to the HNSC scenario. On further examination, we found that there was significant up-regulation of neutrophils as well as the cytokine, IL33 in mutant-*CASP8* HNSCs, both of which were not observed in mutant-*CASP8* UCECs.

**Conclusions:** These results suggested that carcinomas with mutant *CASP8* have broadly similar immune signatures albeit with different effects on survival. We hypothesize that subtle tissue-dependent differences could influence survival by modifying the micro-environment of mutant-*CASP8* carcinomas. High neutrophil numbers, which is a well-known negative prognosticator in HNSCs, and/or high IL33 levels may be some of the factors affecting survival of mutant-*CASP8* cases.

## Background

Exome sequencing, RNA-sequencing, and copy number variation analysis of different cancers have revealed a cornucopia of disease-relevant mutations and altered pathways. The identified genes included those with broad relevance across different cancers, as well as genes that were relevant only in one or a few cancer types. Arguably, the next phase will involve parsing this voluminous data to generate ideas and hypotheses with the potential for clinical impact, and then testing them experimentally.

We are particularly interested in the heterogeneous group of Head and Neck Squamous cell Carcinomas (HNSCs) as these account for a large number of mortalities each year in the Indian subcontinent. Multiple exome sequencing studies have revealed the landscape of recurrent somatic mutations in HNSCs and its prevalent subtype of Oral Squamous Cell Carcinomas (OSCCs) [1–5]. While *TP53* was the most significant recurrently mutated gene in this cancer type, several other genes such as *CASP8, FAT1*, and *NOTCH1* were also unearthed as significantly recurrently mutated by these large-scale sequencing studies. The precise roles of these genes in oral epithelium homeostasis, and how this is altered owing to their mutation in cancer remain to be fully elucidated [6]. In this study, we chose to focus on the *CASP8* gene, which is mutated in 34% of cases with OSCC of the gingiva-buccal sulcus (OSCC-GB), and in ~10% of HNSC patients [1,2,4].

Interestingly, in addition to HNSC, *CASP8* is also recurrently mutated in about 10% of Uterine Corpus Endometrial Carcinoma (UCEC) cases. Here again, the role of *CASP8* in endometrial tissue homeostasis, and how this is altered owing to its mutation in UCEC remains unclear. *CASP8* is also mutated in other cancer types, however, the numbers of such tumors are too low for meaningful analyses. Thus, using the sequencing data on ~500 head and neck and uterine corpus endometrial carcinoma tumors available in The Cancer Genome Atlas (TCGA) [7,8], we sought to identify distinctive features of mutant-*CASP8* tumors.

CASP8 regulates two pathways of programmed cell death; it is a key protease required for the initiation of the extrinsic apoptotic pathway that is targeted by some drug-resistant tumors, and it is an important negative regulator of necroptosis [9–12]. Loss-of-function mutations in *CASP8* could lead to reduced apoptosis and promote tumor survival. It could also lead to enhanced necroptosis and promote tumor cell death. Interestingly, it has been proposed that the necroptotic pathway could be utilized to develop anti-cancer treatments for countering cancers with resistance to apoptosis [13]. On the background of these observations, tumors harboring *CASP8* mutations offer a tractable, physiologically relevant opportunity to understand the changes brought about by *CASP8* mutation, how it affects survival, and if *CASP8* or the necroptotic pathway could be a potential drug target.

In this study, we describe the comparison of RNA-sequencing (RNA-seq) data from head and neck squamous cell carcinoma, and later from uterine corpus endometrial carcinoma, that are mutant or wild type for *CASP8*. We report distinctive molecular features of mutant-*CASP8* HNSCs and UCECs that this comparison revealed. In addition, we describe results obtained by correlating these features to overall survival in the affected patients.

## Methods

### Differential Gene Expression Analysis of wild-type-*CASP8* and mutant-*CASP8* cases

Data for 528 head and neck squamous cell carcinoma (HNSC) cases available at The Cancer Genome Atlas (TCGA) were downloaded in May-June 2017 from https://portal.gdc.cancer.gov/. Clinical data files, Mutation Annotation Format (MAF) files, and mRNA quantification files such as HTSeq files (files with number of reads aligning to each protein-coding gene) and FPKM-UQ files (files with number of fragments aligning per kilobase of transcript per million mapped reads normalized to upper quartile) were downloaded. The HPV status of HNSC cases at TCGA has been reported earlier [14], and this data was used to assign HPV-positive and HPV-negative cases. Cases with and without *CASP8* mutation were selected as shown in figure 1. Among selected cases, mRNA quantification data was available for 354 cases with wild-type-*CASP8* (*CASP8*-WT) and 53 cases with mutant-*CASP8* (*CASP8*-MT). Raw reads from HTSeq files for these cases were used for edgeR analysis [15], to identify genes that were differentially expressed between *CASP8*-WT and *CASP8*-MT.

**Figure 1.**
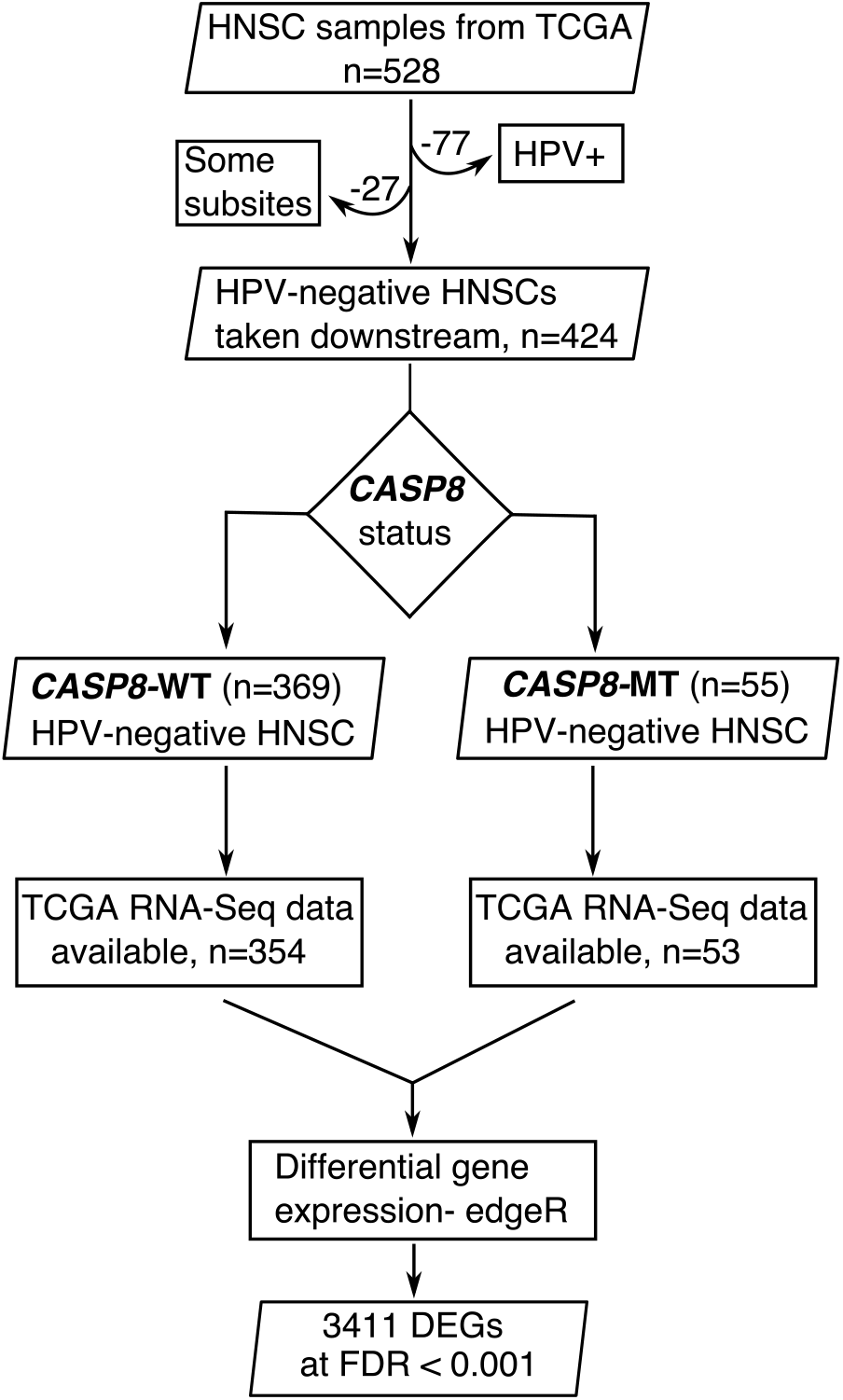
A flowchart indicating the sequence of processes used to select the HNSC cases used in this study. HNSC cases with *CASP8* mutation were identified using MAF files from TCGA. Tumors with *CASP8* mutation occurred in specific sub sites in oral cavity and were HPV-negative. Therefore, HPV-negative HNSC cases with wild-type-CASP8 and the same sub sites were used as control. Gene expression data from HTSeq files of selected cases with *CASP8* mutation and corresponding wild-type control cases was analyzed using edgeR to identify genes that were differentially expressed in *CASP8*-WT or *CASP8*-MT HNSCs. *DEGs:* Differentially Expressed Genes, *FDR:* False Discovery Rate.

Data for 560 uterine corpus endometrial carcinoma (UCEC) cases available at TCGA were downloaded in February 2018. *CASP8* mutations of high or moderate impact were present in 56 UCEC cases. mRNA quantification data was available for 476 *CASP8*-WT tumors and 56 *CASP8*-MT tumors. Raw reads from HTSeq files from these cases were used for edgeR analysis to identify differentially expressed genes. 1231 genes were found to be differentially regulated at FDR<0.001. Of these, 596 genes were downregulated with logFC<1.3 and 258 genes were upregulated with logFC>1.3.

### Gene Ontology and Gene Set Enrichment Analysis (GSEA)

Enrichment analysis to identify biological processes overrepresented among genes differentially expressed between *CASP8*-WT and *CASP8*-MT HNSCs was performed at http://geneontology.org/ [16]. GSEA was performed using a pre-ranked gene list and hallmark gene sets available at the Molecular Signature Database [17]. The hallmark gene sets use either HGNC or entrez gene ids as the gene identifier. Out of the 60,483 transcripts analyzed by edgeR, HGNC gene symbols could be assigned to 37,095. logFC values for these 37,095 genes from the edgeR output from HNSC and UCEC differential gene expression analyses were used to generate the pre-ranked gene list. To perform GSEA with genes that were up-regulated in the skin of mice lacking functional *Caspase-8*, the top 100 up-regulated genes were selected [18]. Of these 100 genes, 80 genes had corresponding human orthologs (*CASP8-* KOSET) as identified using tools available at http://www.informatics.jax.org/. GSEA was then performed using the pre-ranked gene list and the *CASP8*-KOSET.

### Immune Cell Infiltration in HNSC and UCEC cases

Data for distribution of immune infiltrates in TCGA cases was downloaded from Tumor IMmune Estimation Resource (TIMER) at https://cistrome.shinyapps.io/timer/ [19]. Data was available for 353 *CASP8*-WT and 51 *CASP8*-MT cases from HNSC. The comparison of immune cell infiltration levels across *CASP8*-WT, *CASP8*-MT, and HPV-positive cases was performed using a two-sided Wilcoxon rank test and the graphs were plotted using R. Similar analysis was performed for all *CASP8*-MT and *CASP8*-WT cases from UCEC.

### Survival Analysis

Survival analysis was performed to investigate the difference in the survival of *CASP8*-WT and *CASP8*-MT patients from HNSC and UCEC. Survival analysis was also performed to investigate the effect of factors such as expression levels of certain genes and immune cell infiltration on survival. The expression levels of genes of interest were obtained from FPKM-UQ files from TCGA and the data for distribution of immune cell infiltration was obtained from TIMER. Kaplan-Meier curves for *CASP8*-WT and *CASP8*-MT cases were plotted using the Survival and Survminer packages in R and the plots were compared using the log-rank test [20].

To investigate the effect of genes of interest (such as those from gene sets enriched in GSEA or genes involved in necroptosis) and immune cell infiltration levels on survival, multivariate Cox proportional hazards test was performed for HNSC cases. In addition, Cutoff Finder tool available at http://molpath.charite.de/cutoff/ was used to investigate the influence of a single continuous variable on survival [21]. In this analysis of either *CASP8*-WT or *CASP8*-MT cases, the data for each variable was stratified into two groups at the point with the most significant log-rank test.

## Results

### Genes involved in immune response are up-regulated in *mutant-*CASP8** HNSCs

To investigate the significance of *CASP8* mutations in head and neck squamous cell carcinoma (HNSC), we performed differential gene expression analysis using RNA-seq data from HNSC cases with and without *CASP8* mutations, as described in figure 1. *CASP8* mutations were identified using the Mutation Annotation Format (MAF) files. The workflow for somatic mutation calling at TCGA uses four different pipelines: SomaticSniper, MuSE, MuTect2, and VarScan2. Mutation(s) in the *CASP8* gene identified by any one of these four pipelines was included in this study. Using this criterion, *CASP8* mutations were identified in 55 HNSC cases. Notably, the majority (80%) of the identified *CASP8* mutations were predicted by more than one program, and all had either high or moderate impact on function.

As reported previously, HNSCs carrying *CASP8* mutations occurred predominantly in sites within the oral cavity such as the cheek mucosa, floor of mouth, tongue, larynx, and overlapping sites of the lip, oral cavity, and pharynx (Supplementary Table 1). In addition, since HPV-positive (Human Papillomavirus-positive) HNSCs constitute a molecularly distinct subtype; we examined the HPV status of the 55 mutant-*CASP8* HNSCs using data from Chakravarthy *et al* [14]. Based on this reported data, all 55 mutant-*CASP8* HNSCs were found to be HPV-negative. Since all the HNSC cases carrying *CASP8* mutations were HPV-negative, and were from specific sites within the oral cavity, HNSCs carrying wild-type-*CASP8* that were HPV-negative and also from these same sites were selected as controls for all subsequent analyses. A total of 424 HNSC cases of which 369 had wild-type-*CASP8* (*CASP8*-WT) and 55 had mutant-*CASP8* (*CASP8*-MT) were thus selected (Figure 1, see also Supplementary Table 1). Of these, RNA-seq data was available for 354 *CASP8*-WT and 53 *CASP8*-MT cases.

HTSeq files containing raw sequencing reads from *CASP8*-WT and *CASP8*-MT cases were subjected to edgeR analysis for differential gene expression (Supplementary Table 2). At FDR<0.001, 186 genes were up-regulated in *CASP8*-MT with log2FC>1.3 while 1139 genes were down-regulated in *CASP8*-MT with log2FC<–1.3 (Figure 2A). To identify pathways specifically enriched in the *CASP8*-WT or *CASP8*-MT cases, enrichment analysis was performed with the differentially expressed genes using tools available at the Gene Ontology (GO) Consortium. As seen in figure 2A, distinct pathways were enriched in the *CASP8*-WT and *CASP8*-MT cases. Genes involved in the regulation of immune response were enriched in *CASP8*-MT HNSCs while genes with roles in heart development, muscle development, and cellcell signaling were enriched in *CASP8*-WT HNSCs (Supplementary Table 3).

**Figure 2.**
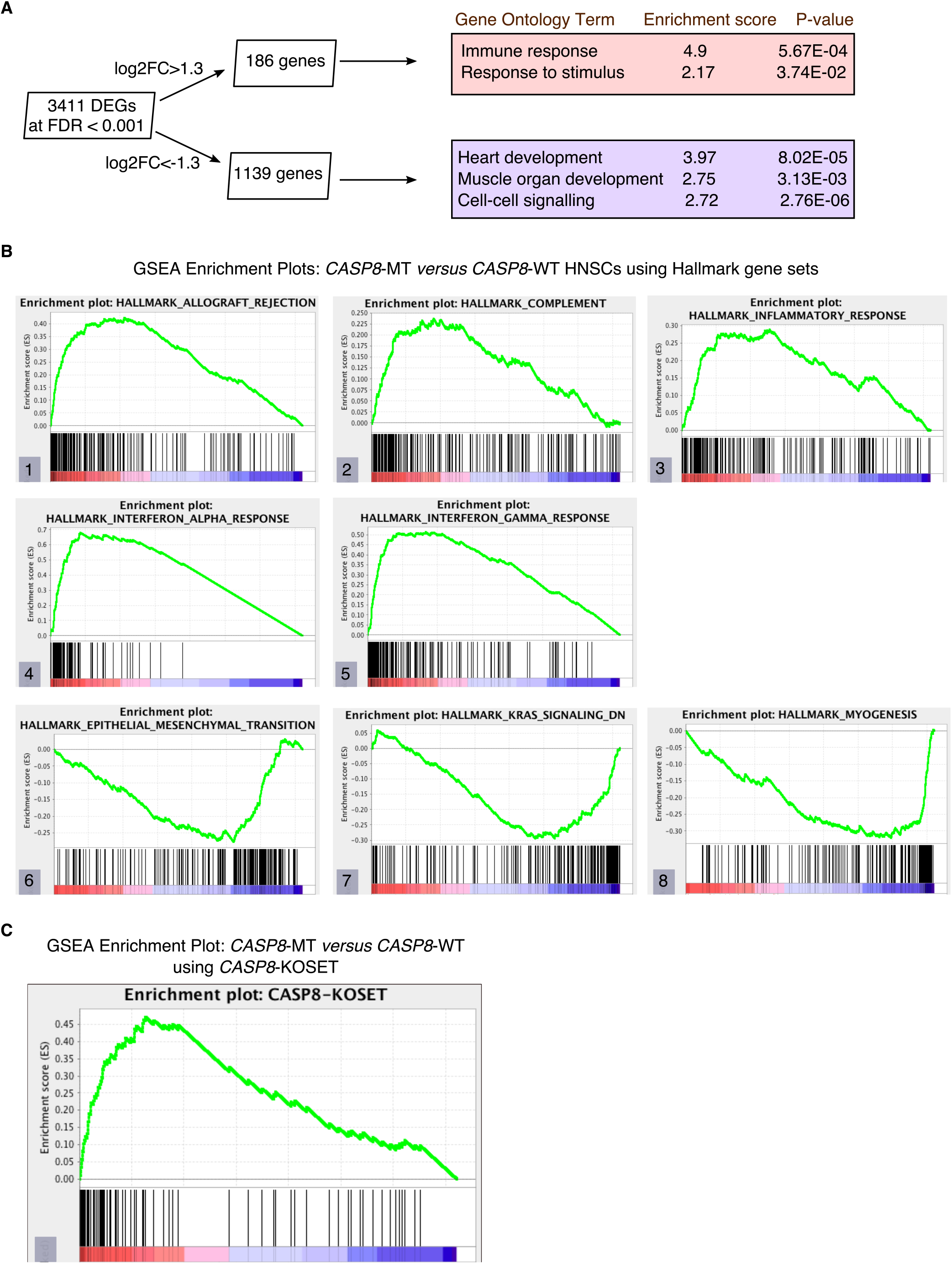
Gene enrichment analyses reveal a prominent immune signature in *CASP8*-MT HNSCs. Gene enrichment analysis was performed using tools available at the Gene Ontology Consortium (A), as well as using the Gene Set Enrichment Analysis tool (B and C). **A**. Enrichment analysis was performed using genes, showing log2FC greater than 1.3 or less than −1.3, and FDR<0.001. The top three gene ontology terms significantly enriched in these gene lists are indicated along with P-values. **B**. GSEA was performed using a pre-ranked list generated using log2FC values from the edgeR analysis. GSEA Hallmark gene sets enriched in *CASP8*-MT HNSCs (plots 1 to 5) or *CASP8*-WT HNSCs are shown (plots 6 to 8). **C**. Enrichment plot of a GSEA performed with the pre-ranked list in panel B and a gene set of human orthologs of the genes up regulated in the skin epidermis of *Casp-8^F/−^K5-Cre* mice (*CASP8*-KOSET) is shown.

We further analyzed the differential gene expression data using the Gene Set Enrichment Analysis (GSEA) tool. After generating a pre-ranked gene list based on logFC values from the edgeR analysis, we queried this list in the GSEA software using hallmark gene sets available at the Molecular Signatures Database. Several gene sets involved in immune regulation such as allograft rejection, complement, inflammatory response, interferon-α response, and interferon-γ response, were specifically enriched in the *CASP8*-MT HNSCs, in sync with the GO results (Figure 2B and Supplementary Table 4). The hallmark gene sets enriched in *CASP8*-WT HNSCs were epithelial-mesenchymal transition (EMT), myogenesis, and the KRAS pathway (Figure 2B and Supplementary Table 4).

### Gene expression in the skins of epidermal ***Caspase-8*** knockout mice mirrors the expression pattern of mutant-*CASP8* HNSCs

Expression of an enzymatically inactive *Caspase-8* mutant or the deletion of wild-type *Caspase-8* in the mouse epidermis leads to chronic skin inflammation [18,22]. A microarray analysis performed by Kovalenko *et al*. to identify genes specifically up-regulated in the skin epidermis of *Casp-8^F/−^K5-Cre* (relative to *Casp-8^F/+^K5-Cre* epidermis) mice revealed increased expression of several immune-regulatory and inflammatory genes including several cytokines. Using the human orthologs of these up-regulated genes (Supplementary Table 5), we again queried the pre-ranked gene list with the GSEA tool. As seen in figure 2C, genes highly expressed in the *Casp-8^F/−^K5-Cre* mouse skins were also significantly enriched in *CASP8*-MT HNSCs (as opposed to their wild-type counterparts), indicating that the inactivation of *CASP8* leads to the up-regulation of a similar set of genes in both mouse and human epidermal tissues.

### Enrichment of immune response gene sets correlates with increased infiltration of specific immune cell types in mutant-*CASP8* HNSCs

Since genes involved in the immune response were specifically enriched in *CASP8*-MT HNSCs, we investigated if this enrichment correlated with the presence or infiltration of immune cells in these tumors. Using the data available at Tumor IMmune Estimation Resource (TIMER) [19], a comprehensive resource for immune cell infiltration of TCGA tumors, we checked if there was a difference in the numbers/types of tumor-associated immune cells between the *CASP8*-WT and *CASP8*-MT HNSCs. Consistent with the GSEA results, *CASP8*-MT cases showed significantly higher infiltration of CD8^+^ T cells, neutrophils, and dendritic cells as compared to *CASP8*-WT cases (p-values<0.0005), suggesting that the immune response to the tumor in WT and MT cases was different (Figure 3A). It has been widely reported that HPV-positive HNSCs have higher infiltration of immune cells as compared to HPV-negative HNSCs [23,24]. In agreement with these reports, HPV-positive HNSCs had significantly higher infiltration of CD8^+^ T cells, neutrophils, and dendritic cells as compared to the *CASP8*-WT HNSCs, but not when compared to the *CASP8*-MT HNSCs (Figure 3A). However, HPV-positive HNSCs also had high levels of infiltration of CD4^+^ T cells and B cells which was not observed in either *CASP8*-WT or *CASP8*-MT HNSCs (Figure 3B).

**Figure 3.**
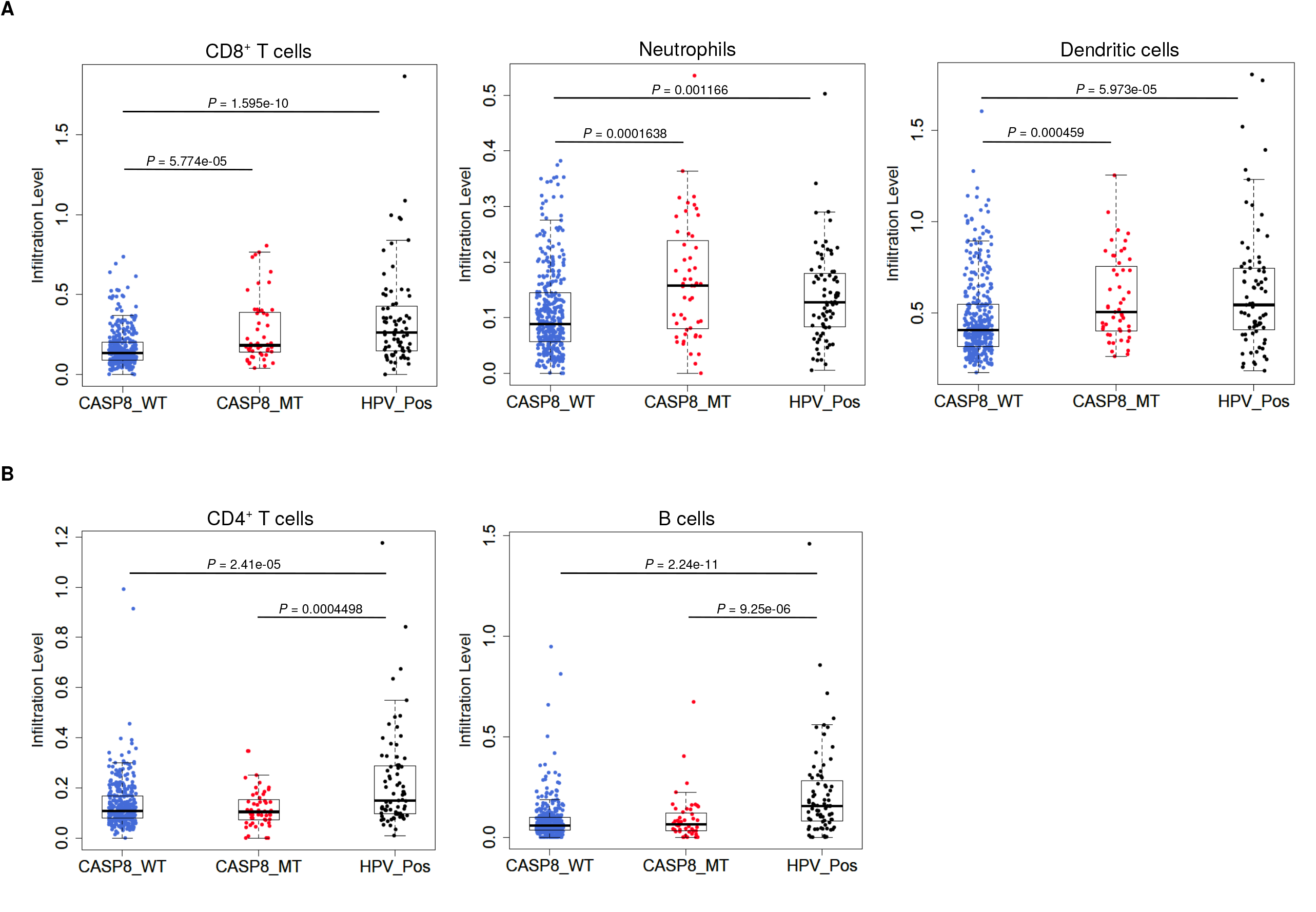
*CASP8*-MT HNSCs have higher numbers of certain types of infiltrating immune cells compared to *CASP8*-WT HNSCs. Immune cell infiltration levels in *CASP8*-WT (blue-filled circles), *CASP8*-MT (red-filled circles), and HPV-positive (black-filled circles) HNSCs were compared using the immune cell infiltration data available at TIMER. Boxplots showing the levels of CD8^+^ T cells, neutrophils, and dendritic cells (A), as well as CD4^+^ T cells and B cells (B) in the three HNSC subsets are displayed. Significance testing was performed using the unpaired two-sided Wilcoxon test. All comparisons with P-value<0.005 were considered significant and are indicated in the plots.

### The “immune signature” of *mutant-*CASP8** HNSCs does not correlate to improved overall survival

High levels of immune cell infiltration in HPV-positive cases correlates with better survival in HPV-positive HNSC cases [23,24]. To investigate if a similar effect could be observed in the survival of HNSC patients with and without *CASP8* mutation, Kaplan-Meier analysis was performed on the *CASP8*-WT and *CASP8*-MT cases (filtered as per the schema in figure 1). In contrast to observations from HPV-positive tumors, there was no significant difference in the survival of patients with and without *CASP8* mutations (p-value=0.16, Figure 4A).

The effect of genes from pathways enriched either in *CASP8*-WT or *CASP8*-MT tumors (listed in Supplementary table 4) on survival was then investigated using the Cox proportional hazards model. Four genes from pathways enriched in *CASP8*-MT HNSCs; *PRF1, CXCR6, CD3D*, and *GZMB*, reduced the hazard ratio significantly in *CASP8*-WT cases but not in *CASP8*-MT cases (Supplementary Table 6). We also performed the survival analysis using Cutoff Finder to investigate the effect of the expression of individual genes on the survival of *CASP8*-WT and *CASP8*-MT cases. Increased expression of all the four genes was significantly associated with higher overall survival in *CASP8*-WT but less so in *CASP8*-MT cases (Figure 4B and Supplementary Figure 1). Similarly, higher CD8^+^ T cell estimates (from TIMER) was also significantly associated with better survival in *CASP8*-WT but not in *CASP8*-MT HNSCs (Figure 4B). Since CASP8 is a negative regulator of the necroptotic pathway [11,12], we also investigated the effect of expression levels of genes involved in necroptosis on survival. Higher expression of *RIPK1, RIPK3*, and *MLKL* was associated with higher overall survival in *CASP8*-WT but not in *CASP8*-MT cases (Supplementary Figure 2). Additional factors that influenced survival are shown in supplementary table 6.

**Figure 4.**
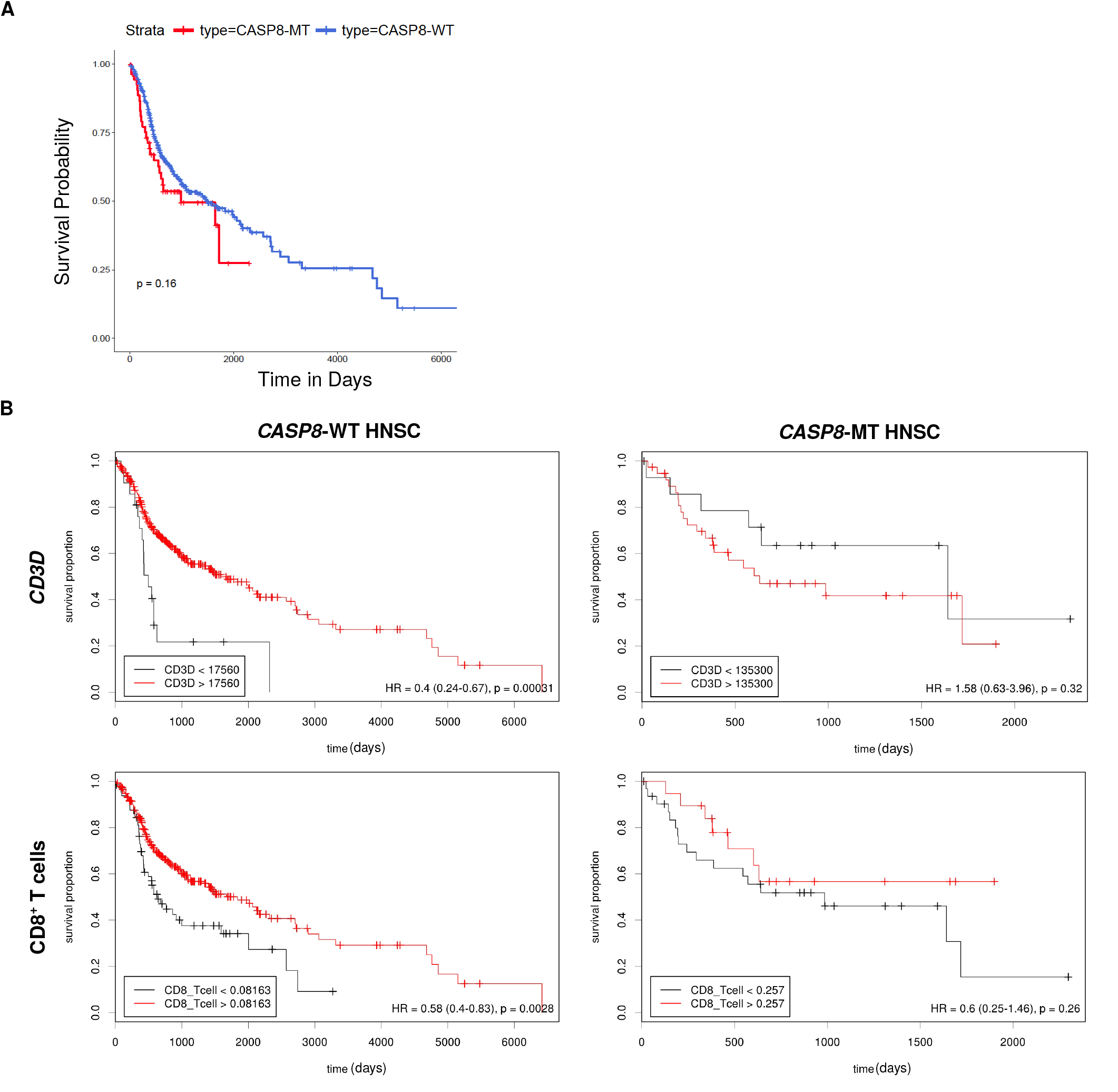
Survival analysis indicates lack of a survival advantage in *CASP8*-MT HNSCs in spite of their immune signature. **A**. Kaplan-Meier plots showing the survival probability of patients with *CASP8*-WT or *CASP8*-MT HNSC tumors (filtered as per the schema in figure 1). Log-rank test was used to compare the two curves and the log-rank P-value is indicated. **B**. Survival plots generated using the Cutoff Finder tool showing the influence of the expression levels of *CD3D* and the levels of CD8^+^ T cells on overall survival in *CASP8*-WT (left) and *CASP8*-MT (right) cases. Gene expression data was obtained from FPKM-UQ files at TCGA and immune cell infiltration data was obtained from TIMER.

### mutant-*CASP8* UCECs exhibit an immune signature similar to mutant-*CASP8* HNSCs

We then investigated if this effect seen in *CASP8*-MT HNSCs was broadly applicable across other cancers carrying *CASP8* mutations. Apart from HNSC, among the cancers covered by the TCGA program, UCECs carried the most numbers of mutations in the *CASP8* gene. About ~10% UCECs harbor *CASP8* mutations. From a total of 560 UCEC cases, RNA-seq data was available for 476 *CASP8*-WT and 56 *CASP8*-MT cases. HTSeq files containing raw sequencing reads from these two groups were subjected to edgeR analysis for differential gene expression and further analyzed using the GSEA tool. Several gene sets involved in immune regulation were specifically enriched in the *CASP8*-MT UCECs. Notably, categories such as allograft rejection, interferon-α response, and interferon-γ response, were enriched in the *CASP8*-MT UCECs similar to the HNSC results. However, unlike HNSCs, the gene set for inflammatory response did not show any enrichment in *CASP8*-MT UCECs. *CASP8*-MT UCECs were additionally enriched for genes involved in apoptosis. Notably, this was not observed in the *CASP8*-MT HNSCs (Figure 5A, see also figure 2B).

**Figure 5.**
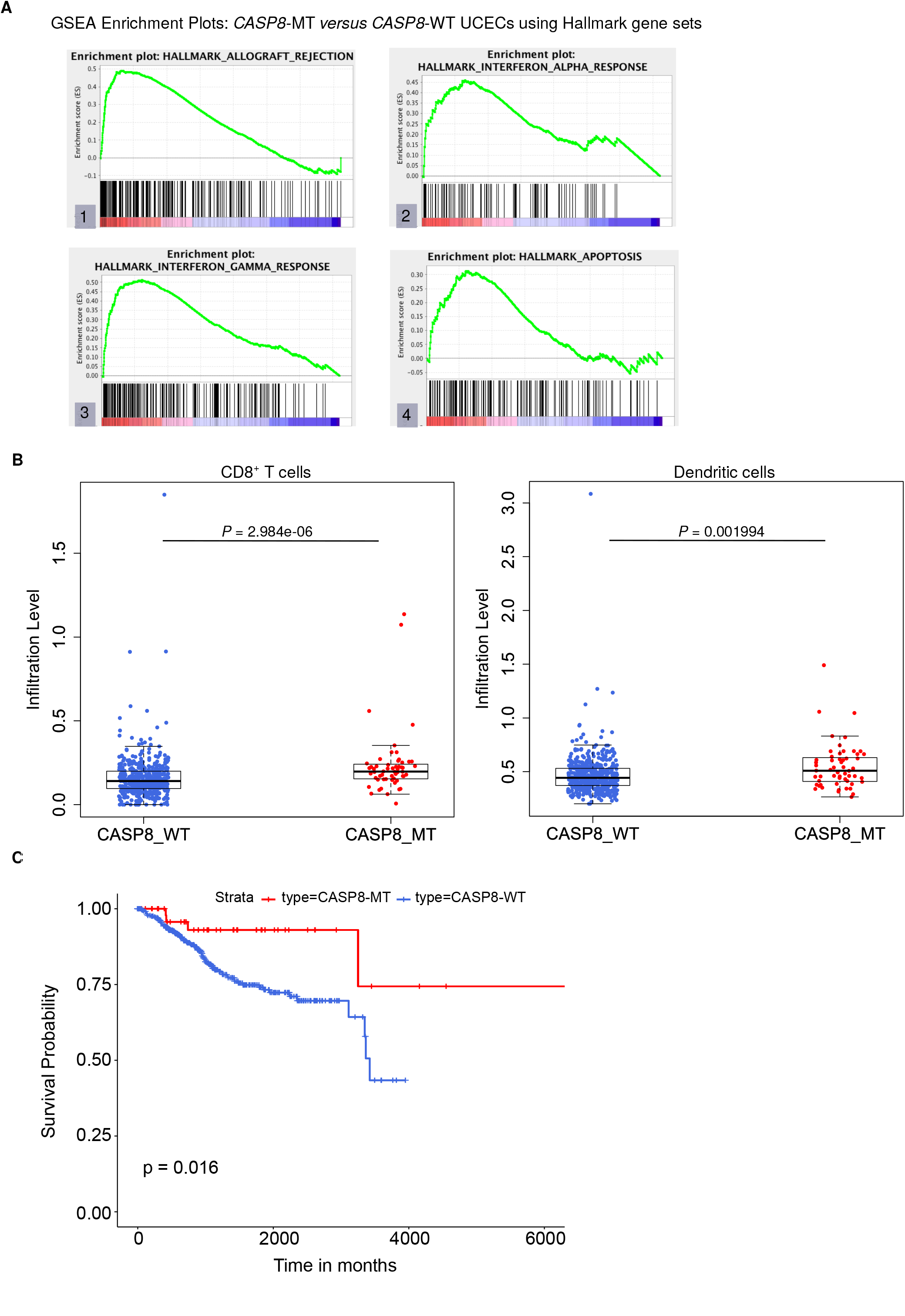
*CASP8*-MT UCECs display an immune gene signature, have higher numbers of certain types of infiltrating immune cells, and survive better than *CASP8*-WT UCECs. **A**. GSEA was performed using a pre-ranked list generated using log2FC values from the edgeR analysis. Some GSEA Hallmark gene sets enriched in *CASP8*-MT UCECs (plots 1 to 5) are shown. **B**. Immune cell infiltration levels in *CASP8*-WT (blue-filled circles) and *CASP8*-MT (red-filled circles) UCECs were compared using the immune cell infiltration data available at TIMER. Boxplots showing the levels of CD8^+^ T cells and dendritic cells in the two UCEC groupsare displayed. Significance testing was performed using the unpaired two-sided Wilcoxon test. All comparisons with P-value<0.005 were considered significant and are indicated in the plots. **C**. Kaplan-Meier plots showing the survival probability of patients with *CASP8*-WT or *CASP8*-MT UCEC tumors. Log-rank test was used to compare the two curves and the log-rank P-value is indicated.

### High levels of IL33 and neutrophil infiltration are observed in mutant-*CASP8* HNSCs but not in mutant-*CASP8* UCECs

Using TIMER, we then checked the levels of infiltrating immune cells in the *CASP8*-WT and *CASP8*-MT UCECs. Consistent with the GSEA results, *CASP8*-MT UCEC cases showed significantly higher infiltration of CD8^+^ T cells and dendritic cells as compared to *CASP8*-WT cases (p-values<0.005). However, in contrast to the HNSC data, the levels of neutrophils were not significantly higher in the *CASP8*-MT UCEC group (Figure 5B, see also figure 3A). We then investigated if differences in the levels of neutrophil-active chemokines could potentially explain this observation [25]. From the edgeR differential expression data comparing the *CASP8*-MT and *CASP8*-WT groups in HNSC and UCEC, we obtained the fold change values and statistical significance of different chemokines known to attract neutrophils (Supplementary table 7). Interestingly, the cytokine IL33 was significantly up regulated (1.8 fold, FDR<0.001) in *CASP8*-MT HNSCs but not in *CASP8*-MT UCECs. Next, we performed Kaplan-Meier analysis on *CASP8*-WT and *CASP8*-MT UCEC cases. In contrast to the HNSC survival data, there was a difference in the survival of UCEC cases with and without *CASP8* mutations, with cases harboring *CASP8* mutations reporting better overall survival (p-value=0.019, Figure 5C).

**Figure 6.**
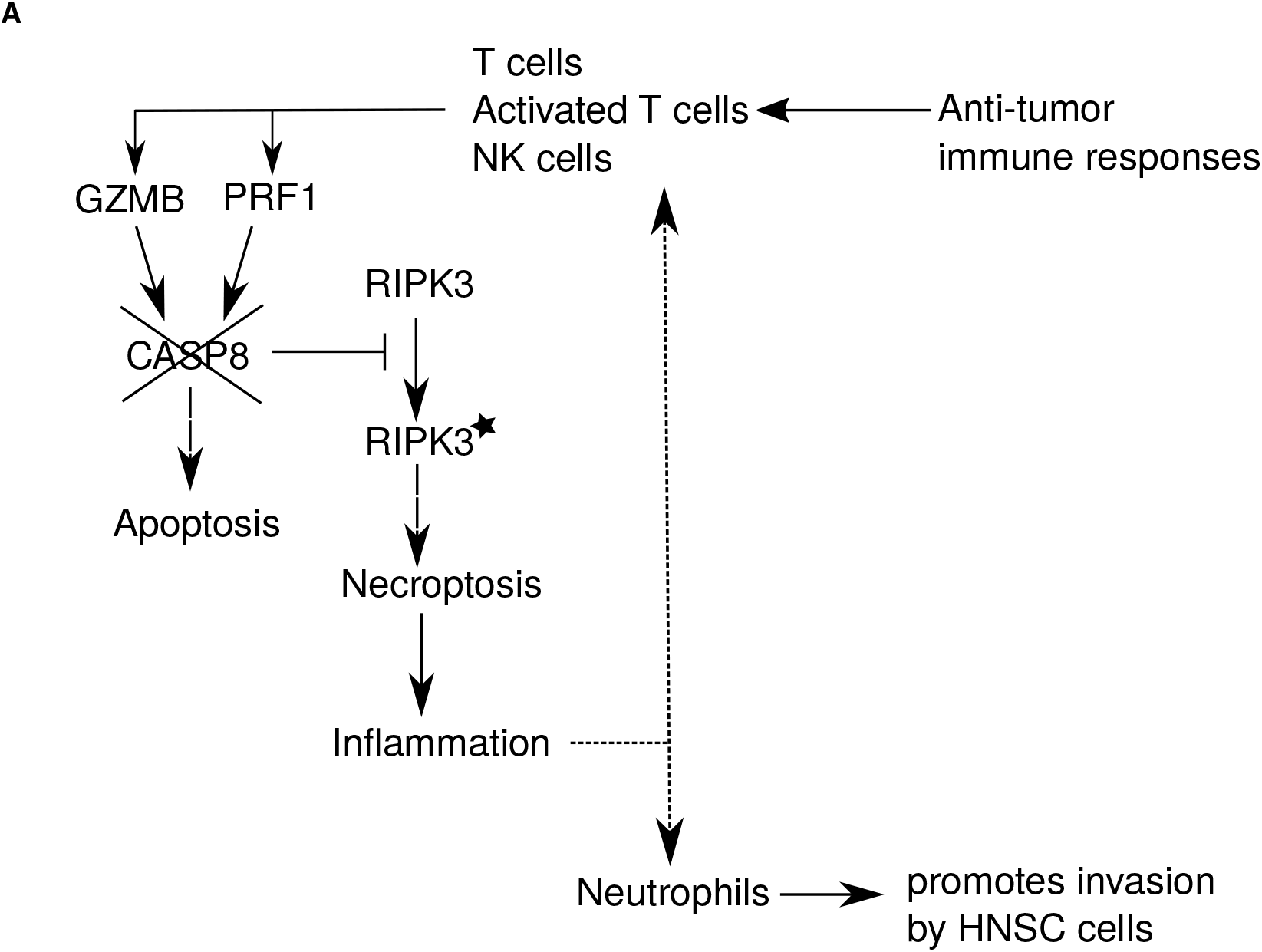
Model for significance of *CASP8* mutation in HNSC. Straight lines represent known pathways, and dotted lines represent possible effects of *CASP8* mutation. Immune response to HNSC leads to T cell activation followed by apoptosis of tumor cells. The expression of genes such as *PRF1* and *GZMB* required for apoptosis facilitates this process. Tumors with mutation in *CASP8* are protected from apoptosis and such patients lose the survival advantage associated with T cell activation and high expression of apoptosis-related genes. *CASP8* mutation also promotes necroptosis that leads to further inflammation, production of alarmins such as IL33, and infiltration of certain immune cells including neutrophils that are known to promote invasion in OSCC. However, unlike other pro-inflammatory situations such as in HPV-positive tumors, the particular inflammatory microenvironment created due to *CASP8* mutation does not appear to provide survival advantage to the patients.

## Discussion

Here, we report a distinct class of carcinomas that have mutated *CASP8*. Using bioinformatics approaches to mine the TCGA data, we identified immune response-related genes as a prominent shared category of genes enriched in *CASP8*-MT carcinomas. We then explored the correlation of this gene set enrichment with survival. Our analyses showed that despite similarities in the enrichment of gene sets, these carcinomas exhibited varying correlations of immune signature with survival. In the first part of our analyses, we investigated the implications of the enrichment of this immune signature across different HNSC subtypes. Subsequently, in the second part, we investigated the correlation between immune signature and survival in two carcinomas having a significant number of cases with *CASP8* mutations, HNSC and UCEC. Our studies indicated that tissue-specific differences, such as the levels of infiltrating neutrophils and the cytokine IL33, could be responsible for the varying correlation of immune signature with survival.

Multiple studies have reported that HPV-positive HNSCs display a strong immune signature and high infiltration of immune cells that correlates with better survival. In contrast, our studies show that the enrichment of immune response genes and infiltration of immune cells in *CASP8*-MT HNSCs does not correlate with improved prognosis. In fact, *CASP8* mutation leads to the loss of a survival advantage that is observed in HNSC patients with wild-type *CASP8* tumors under certain conditions. These results argue that a tumor microenvironment with high infiltration of immune cells does not necessarily provide a survival benefit in HNSCs.

We can think of at least two potential scenarios to explain the increased immune cell infiltration observed in *CASP8*-MT tumors. (a) Unregulated inflammatory and wound healing response: As mentioned earlier, loss of *Caspase-8* in the mouse epidermis leads to chronic inflammation [18]. The infiltration of immune cells in mucosa lacking *CASP8* accompanied by the enrichment of immune-associated gene sets is highly reminiscent of this phenotype. It has also been proposed that the loss of *Caspase-8* in the mouse skin epidermis simulates a wound healing response [22]. Both scenarios involve a gamut of immune cell types and secreted cytokine factors, leading to immune cell infiltration. It should however be noted that although similar gene sets are enriched in mouse skins lacking *Caspase-8* and in *CASP8*-MT tumors, the types of immune cell infiltrates in the two are different. (b) Necroptosis: More recently, several studies have revealed a role for Caspase-8 as an inhibitor of necroptosis, a highly pro-inflammatory mode of cell death [9,10]. In intestinal epithelia, the loss of *Caspase*-8 promoted necroptosis through the activation of RIP kinases and MLKL [11,12]. A similar scenario could be occurring in *CASP8*-MT tumors leading to the expression of pro-inflammatory genes and the infiltration of immune cells (see model in figure 6).

Why doesn’t the increased number of immune cells translate into improved prognosis in *CASP8*-MT HNSC tumors? Since CASP8 is an important mediator of the extrinsic apoptotic pathway, *CASP8*-MT tumors may have greater resistance to Fas- or DR5- mediated cell death pathways, which are typically employed by CD8^+^ T cells and Natural Killer cells to target infected/tumor cells [26,27]. The survival analysis carried out in this study showed that *CASP8*-WT HNSC patients with higher expression of genes involved in T-cell mediated cytotoxicity had better survival. Importantly, this advantage was not seen in *CASP8*-MT patients.

Several studies have reported that high neutrophil numbers and an elevated neutrophil/lymphocyte ratio portended poorer prognosis in OSCC [28,29]. Thus, it is possible that elevated levels of neutrophil infiltration seen in *CASP8*-MT HNSC cases could be one of perhaps several events contributing to the poorer prognosis of *CASP8*-MT HNSCs (see model in figure 6). IL33, a cytokine and an alarmin linked to necroptosis may represent a possible mechanism for neutrophil recruitment in these cases [30,31]. High IL33 levels are also associated with poor prognosis in HNSCs [32]. In addition, the pro-inflammatory environment generated during necroptosis may hold other advantages for the survival of *CASP8*-MT HNSCs. Necroptosis, IL33 levels, and neutrophil infiltration together or through independent mechanisms could be leading to a pro-tumor environment. Thus, promoting necroptosis may not necessarily translate into better survival for HNSC patients with apoptosis-resistant tumors.

Another reason for the lack of survival advantage in *CASP8*-MT HNSCs could be the composition of tumor-infiltrating immune cells in these tumors. For instance, HPV-positive tumors had higher levels of B cells and CD4^+^ T cells as compared to *CASP8*-MT tumors. It is likely that in addition to cytotoxic T cells, B cells and CD4^+^ T cells are required to mediate an immune response essential for tumor cell death, possibly for tumor antigen presentation or cytokine secretion.

A comparison of *CASP8*-MT HNSCs and *CASP8*-MT UCECs highlighted important differences between the two carcinomas. Notably, *CASP8*-MT UCECs showed a significant survival advantage over *CASP8*-WT UCECs, unlike its HNSC counterpart. While we do not yet know the causal reason(s), the differences *per se* may be worth noting and could be responsible for this advantage. For instance, in contrast to *CASP8*-MT HNSCs, the gene set for inflammatory response was not enriched but the gene set for apoptosis was enriched in *CASP8*-MT UCECs. There was also no increased infiltration of neutrophils or transcriptional upregulation of IL33 in *CASP8*-MT UCECs. The up-regulation of apoptotic pathways together with the lack of enrichment of an inflammation-associated gene set that is typical of necroptosis perhaps indicates that apoptosis, rather than necroptosis, is the predominant mode of programmed cell death in *CASP8*-MT UCECs. This lack of inflammation may also be responsible for the lack of neutrophil infiltration in *CASP8*-MT UCECs since neutrophil chemoattractants, such as IL33, may not be released during apoptosis but is perhaps released during the highly inflammatory process of necroptosis, in turn leading to neutrophil infiltration.

It is also possible that necroptosis is initiated in *CASP8*-MT UCECs but the accompanying IL33 up-regulation and/or neutrophil infiltration seen in HNSCs does not take place due to tissue-specific differences. Under such conditions, apoptosis and necroptosis together could provide the survival advantage that is observed in *CASP8*-MT UCECs. Thus, in contrast to HNSCs, Caspase-8 pathway can be explored to identify potential drug targets in UCECs.

## Conclusions

In this *in silico* study, we explore the functional implications of *CASP8* mutations that have been identified across carcinomas through large-scale genomic studies. Our studies show that CASP8-mutated carcinomas display a shared immune signature. However, the consequences of this immune signature vary with *CASP8*-MT UCECs showing better survival while *CASP8*-MT HNSC cases do not have any survival advantage. Our analyses indicate that these differences could perhaps be attributed to differences in neutrophil infiltration and/or IL33 levels in these carcinomas. Furthermore, it highlights the need to further investigate the interaction between pathways of programmed cell death, immune response, and survival in carcinomas. Such studies could open a new window for therapeutic intervention in *CASP8*-mutated carcinomas.

## Authors’ contributions

YG and SS contributed equally to the conception and design of the study, and the analysis and interpretation of data. The authors jointly wrote the manuscript and prepared the figures. Both the authors have approved of the version submitted for publication.

## Acknowledgements

The results shown here are based upon data generated by the TCGA Research Network: http://cancergenome.nih.gov/. We thank patients who donated samples to the TCGA and consented to share the resulting data. We also wish to thank TCGA for the unrestricted access provided to the data used in this work. We would like to thank Amrendra Mishra (Hannover Biomedical Research School), Urvashi Bahadur (Strand Life Sciences), and Colin Jamora (inStem) for their helpful comments on this manuscript.

## References

1. Agrawal N, Frederick MJ, Pickering CR, Bettegowda C, Chang K, Li RJ, et al. Exome Sequencing of Head and Neck Squamous Cell Carcinoma Reveals Inactivating Mutations in NOTCH1. Science 2011; 333:1154–7.

2. Maitra A, Biswas NK, Amin K, Kowtal P, Kumar S, Das S, et al. Mutational landscape of gingivo-buccal oral squamous cell carcinoma reveals new recurrently-mutated genes and molecular subgroups. Nat Commun. 2013; 4.

3. Pickering CR, Zhang J, Yoo SY, Bengtsson L, Moorthy S, Neskey DM, et al. Integrative genomic characterization of oral squamous cell carcinoma identifies frequent somatic drivers. Cancer Discov. 2013; 3:770–81.

4. Stransky N, Egloff AM, Tward AD, Kostic AD, Cibulskis K, Sivachenko A, et al. The Mutational Landscape of Head and Neck Squamous Cell Carcinoma. Science 2011; 333:1157–60.

5. Hayes TF, Benaich N, Goldie SJ, Sipilä K, Ames-Draycott A, Cai W, et al. Integrative genomic and functional analysis of human oral squamous cell carcinoma cell lines reveals synergistic effects of FAT1 and CASP8 inactivation. Cancer Lett. 2016; 383:106–14.

6. Rothenberg SM, Ellisen LW. The molecular pathogenesis of head and neck squamous cell carcinoma. J Clin Invest. 2012; 122:1951–7.

7. Lawrence MS, Sougnez C, Lichtenstein L, Cibulskis K, Lander E, Gabriel SB, et al. Comprehensive genomic characterization of head and neck squamous cell carcinomas. Nature 2015; 517:576–82.

8. Getz G, Gabriel S, Cibulskis K, Lander E, Sivachenko A, Sougnez C, et al. Integrated genomic characterization of endometrial carcinoma. Nature 2013; 497:67–73.

9. Pasparakis M, Vandenabeele P. Necroptosis and its role in inflammation. Nature 2015; 517:311–20.

10. Feltham R, Vince JE, Lawlor KE. Caspase-8: not so silently deadly. Clin Transl Immunol. 2017; 6:e124.

11. Günther C, Martini E, Wittkopf N, Amann K, Weigmann B, Neumann H, et al. Caspase-8 regulates TNF-α-induced epithelial necroptosis and terminal ileitis. Nature 2011; 477:335–9.

12. Weinlich R, Oberst A, Dillon CP, Janke LJ, Milasta S, Lukens JR, et al. Protective Roles for Caspase-8 and cFLIP in Adult Homeostasis. Cell Rep. 2013; 5:340–8.

13. Su Z, Yang Z, Xie L, DeWitt JP, Chen Y. Cancer therapy in the necroptosis era. Cell Death Differ. 2016; 23:748–56.

14. Chakravarthy A, Henderson S, Thirdborough SM, Ottensmeier CH, Su X, Lechner M, et al. Human papillomavirus drives tumor development throughout the head and neck: Improved prognosis is associated with an immune response largely restricted to the Oropharynx. J Clin Oncol. 2016; 34:4132–41.

15. Robinson MD, McCarthy DJ, Smyth GK. edgeR: a Bioconductor package for differential expression analysis of digital gene expression data. Bioinformatics 2010; 26:139–40.

16. The Gene Ontology Consortium. Expansion of the Gene Ontology knowledgebase and resources. Nucleic Acids Res. 2017; 45:D331–8.

17. Subramanian A, Tamayo P, Mootha VK, Mukherjee S, Ebert BL, Gillette MA, et al. Gene set enrichment analysis: a knowledge-based approach for interpreting genome-wide expression profiles. Proc Natl Acad Sci U S A. 2005; 102:15545–50.

18. Kovalenko A, Kim J-C, Kang T-B, Rajput A, Bogdanov K, Dittrich-Breiholz O, et al. Caspase-8 deficiency in epidermal keratinocytes triggers an inflammatory skin disease. J Exp Med. 2009; 206:2161–77.

19. Li T, Fan J, Wang B, Traugh N, Chen Q, Liu JS, et al. TIMER: A Web Server for Comprehensive Analysis of Tumor-Infiltrating Immune Cells. Cancer Res. 2017; 77:e108–10.

20. Therneau TM. Survival Analysis [R package survival version 2.41-3] [Internet]. Comprehensive R Archive Network (CRAN); [cited 2017 Nov 21]. Available from: https://cran.r-project.org/web/packages/survival/index.html

21. Budczies J, Klauschen F, Sinn B V., Győrffy B, Schmitt WD, Darb-Esfahani S, et al. Cutoff Finder: A Comprehensive and Straightforward Web Application Enabling Rapid Biomarker Cutoff Optimization. van Diest P, editor. PLoS One 2012; 7:e51862.

22. Lee P, Lee D-J, Chan C, Chen S-W, Ch’en I, Jamora C. Dynamic expression of epidermal caspase 8 simulates a wound healing response. Nature 2009; 458:519–23.

23. Nguyen N, Bellile E, Thomas D, McHugh J, Rozek L, Virani S, et al. Tumor infiltrating lymphocytes and survival in patients with head and neck squamous cell carcinoma. Head Neck. 2016; 38:1074–84.

24. Russell S, Angell T, Lechner M, Liebertz D, Correa A, Sinha U, et al. Immune cell infiltration patterns and survival in head and neck squamous cell carcinoma. Head Neck Oncol. 2013; 5:24.

25. Sadik C, Kim N, Luster A. Neutrophils cascading their way to inflammation. Trends in Immunology 2011,32:452–460.

26. Li C, Egloff AM, Sen M, Grandis JR, Johnson DE. Caspase-8 mutations in head and neck cancer confer resistance to death receptor-mediated apoptosis and enhance migration, invasion, and tumor growth. Mol Oncol 2014; 8:1220–30.

27. Rooney MS, Shukla SA, Wu CJ, Getz G, Hacohen N. Molecular and genetic properties of tumors associated with local immune cytolytic activity. Cell 2015; 160:48–61.

28. Mahalakshmi R, Boaz K, Srikant N, Baliga M, Shetty P, Prasad M, et al. Neutrophil-to-lymphocyte ratio: A surrogate marker for prognosis of oral squamous cell carcinoma. Indian J Med Paediatr Oncol 2018; 39:8–12.

29. Glogauer JE, Sun CX, Bradley G, Magalhaes MA. Neutrophils increase oral squamous cell carcinoma invasion through an invadopodia-dependent pathway. Cancer Immunol Res. 2015; 3:1218–26.

30. Alves-Filho JC, Sônego F, Souto FO, Freitas A, Verri WA Jr, Auxiliadora-Martins M, et al. Interleukin-33 attenuates sepsis by enhancing neutrophil influx to the site of infection. Nat Med. 2010; 16:708–12.

31. Hueber AJ, Alves-Filho JC, Asquith DL, Michels C, Millar NL, Reilly JH, et al. IL-33 induces skin inflammation with mast cell and neutrophil activation. Eur. J. Immunol. 2011; 41:2229–2237.

32. Chen S, Nieh S, Jao S, Wu M, Liu C, Chang Y, et al. The paracrine effect of cancer-associated fibroblast-induced interleukin-33 regulates the invasiveness of head and neck squamous cell carcinoma. J Pathol. 2013; 231:180–189.

